# Minimal condition repetitions required in rapid serial visual presentation decoding paradigms

**DOI:** 10.1101/2023.05.30.542960

**Authors:** Tijl Grootswagers

**Affiliations:** The MARCS Institute for Brain, Behaviour, and Development, Western Sydney University, Sydney, NSW, Australia

## Abstract

Rapid Serial Visual Presentation (RSVP) decoding paradigms allow testing a greater number of conditions than was previously possible within short experimental sessions. However, in these designs individual neural responses may be more susceptible to noise due to responses overlapping with adjacent epochs. This study investigates the minimum number of repetitions required for reliable decoding accuracies in RSVP decoding paradigms. We used previously published EEG data and conducted a standard decoding analysis while varying the number of repetitions used. We found that it is possible to obtain reliable decoding accuracies with only around six repetitions of each condition, which has important implications for research questions that require short experiments, particularly for studying populations who may not be able to tolerate longer or more demanding protocols. These findings highlight the potential benefits of using efficient RSVP decoding designs and conducting short experiments and may have far-reaching impacts in cognitive neuroscience, by providing insights into optimizing data collection methods for diverse populations and experimental protocols.

## Introduction

Recent interest has emerged in combining decoding approaches with rapid serial visual presentation (RSVP) designs as an efficient means of obtaining substantial amounts of neural data. RSVP decoding paradigms have the potential to test a greater number of conditions than was previously possible within short experimental sessions, making these methods highly desirable for visual perception studies (see e.g., Grootswagers et al., 2021, 2022; Shatek et al., 2022) or studies that investigate how the brain processes temporal sequences (Grootswagers et al., 2019; King & Wyart, 2021; Marti & Dehaene, 2017; Mohsenzadeh et al., 2018; Robinson et al., 2019).

However, while RSVP decoding enables the presentation of many stimuli in quick succession, individual neural responses may be more susceptible to noise due to responses overlapping with adjacent epochs, and strong masking effects (Robinson et al., 2019). Therefore, it remains unclear how many repetitions are necessary to obtain dependable decoding results. The aim of the current study was to establish the minimum number of repetitions required in RSVP decoding designs. The resulting lower bound estimates on repetition numbers will be informative for a broad range of future experimental studies.

## Methods

We used previously published electroencephalography (EEG) data which investigated the neural basis of visual object recognition using RSVP decoding (Grootswagers et al., 2019). The original study (n=16) comprised 200 visual objects presented in random order in 40 sequences at 5Hz with a duration of 200ms each. We applied a minimal preprocessing pipeline in EEGlab (Delorme & Makeig, 2004), which consisted of a high-pass filter at 0.1Hz, low-pass filter at 100Hz, and downsampling to 250Hz. To segment the data, we created epochs that were locked to each individual stimulus presentation, ranging from 100 before to 600ms following stimulus onset. Note that these preprocessing steps are the same as those employed in the original study (Grootswagers et al., 2019).

To classify between the 200 different visual stimuli, we conducted a sliding window time-series decoding analysis (Grootswagers et al., 2017), using a leave-one-sequence-out cross-validation scheme and a regularized Linear Discriminant Classifier implemented in the CoSMoMVPA toolbox (Oosterhof et al., 2016). To estimate the minimum number of repetitions required to fit the LDA classifiers, we incrementally added single sequences as each sequence contains one repetition of every stimulus, starting with the first three sequences up to the maximum of 40 sequences. This approach enabled us to simulate having stopped the original experiment earlier, while assessing the robustness and consistency of the results at each stage. All materials to reproduce these results are publicly available: https://github.com/Tijl/RSVP-repetitions-test

## Results

The relationship between the number of repetitions and decoding accuracy are presented in Figure 1. Our results showed that increasing the number of repetitions leads to improved decoding accuracy. We firstly observed a fast increase in decoding accuracy as the number of repetitions increased, with smaller gains after around 15 repetitions (Fig. 1A/D). To assess the reliability of the decoding effects, we applied statistical thresholding to the results (Fig. 1B/C). This analysis revealed that the decoding effects were reliable starting from about six repetitions, and the number of above-chance points increased thereafter.

**Figure 1.**
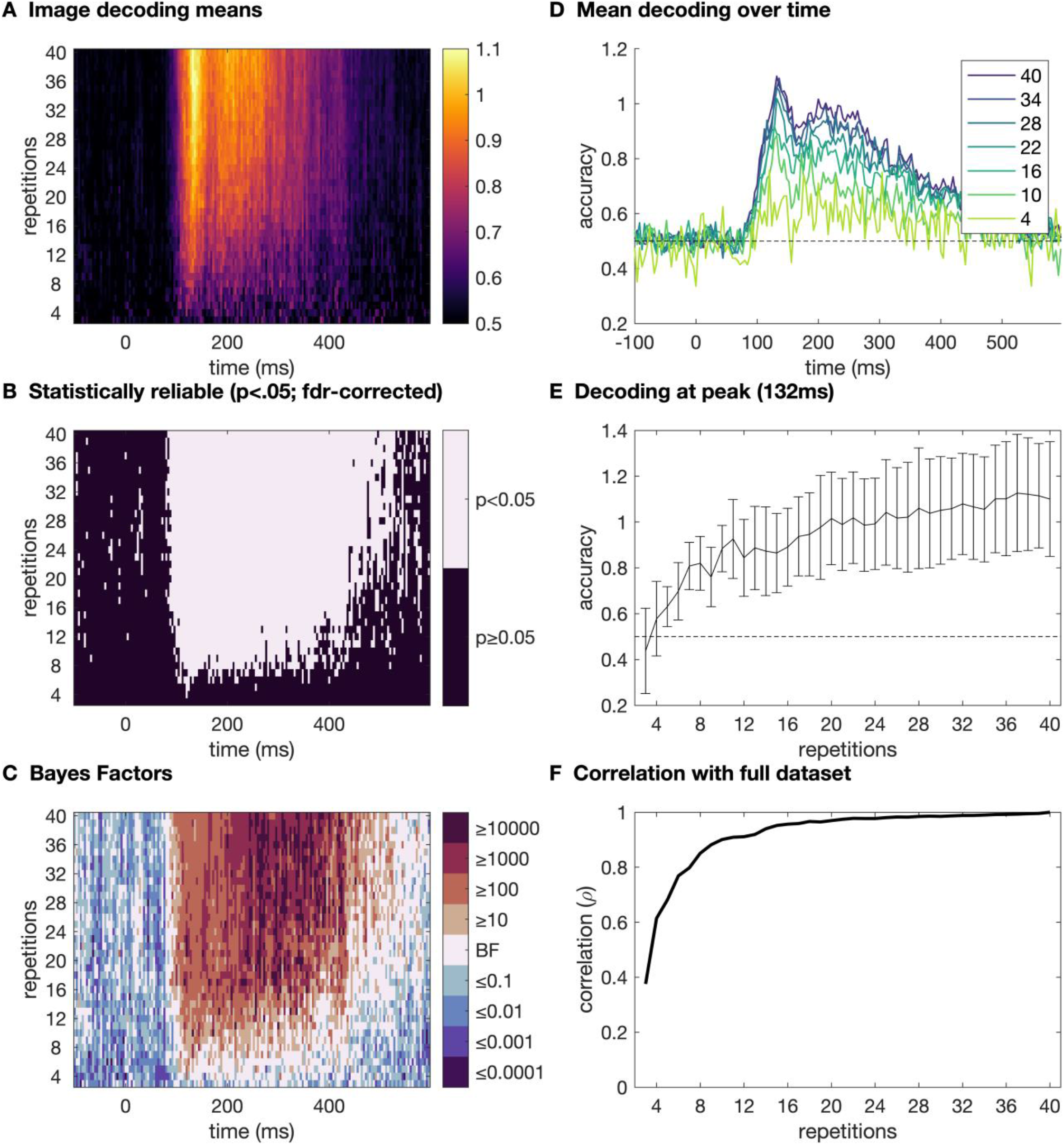
**A**. Time-varying subject-averaged (n=16) decoding accuracies. Brighter colours indicate higher accuracies and rows show the results of different numbers of repetitions. **B**. Statistical thresholding of the results shows where reliable decoding effects are present in the data. **C**. Bayes Factors showing evidence strength for the null and alternative hypothesis of at-chance versus above-chance decoding. **D**. Time-varying subject-averaged decoding accuracies for selected numbers of repeats. Chance-level is 0.5% (200 images). **E**. Comparison of accuracies as a function of the number of repetitions at one time point, including 95% confidence intervals across subjects. Chance-level is 0.5% (200 images). **F**. Correlating the decoding time course obtained at each repetition numbers with the full dataset shows how similar the trajectories are to the complete dataset.

To further explore the relationship between the number of repetitions and decoding accuracy, we compared accuracies as a function of the number of repetitions at the average peak decoding time (Fig. 1E). We found that the decoding accuracy increased as the number of repetitions increased, but that the variance also grew with higher numbers of repetitions.

Finally, to examine the extent to which the decoding trajectories obtained at each repetition increase resembled the complete dataset, we correlated the decoding time course obtained at each number of repetitions with the full dataset (Fig. 1F). We observed a high degree of similarity between the decoding trajectories obtained at each number of repetitions and the full dataset, starting at 0.4 for 3 repetitions, but quickly increasing to 0.77 at 6 repetitions.

## Discussion

This study has revealed that it is possible to obtain reliable decoding accuracies using as little as six repetitions, which in the original study was achieved in about 4 minutes (40 seconds per RSVP sequence). This observation is consistent with similar investigations that were performed on trial numbers in non-RSVP studies (Teichmann et al., 2022). This result has important implications for research questions that require short experiments. This is particularly relevant for studying populations who may not be able to tolerate longer or more demanding protocols, such as in young infants or in ADHD. Our results suggest only five minutes of RSVP data may be sufficient obtain accurate results which highlights RSVP designs as a promising future avenue for studying visual perception these populations.

On the other hand, the low number of required repetitions per stimulus allows for massively scaling up the overall number of stimuli used in a single study. For example, recent work recorded EEG data 12 repetitions of 1854 object concepts in a one-hour study (Grootswagers et al., 2022), and the current results can be used to guide the collection of future large datasets using similar paradigms. Overall, our work highlights the potential benefits of using efficient RSVP decoding designs in large condition-rich, or in short powerful experiments. These findings could have wide-reaching impacts in cognitive neuroscience, by providing guidance into optimizing data collection methods for diverse populations and experimental protocols.

## Acknowledgments

This work was supported by Australian Research Council Grant DE230100380. An earlier version of this manuscript was accepted for presentation at the 2023 Cognitive Computational Neuroscience meeting (https://2023.ccneuro.org/). We thank the 11 anonymous conference reviewers for their suggestions, which are incorporated in the current version.

## Notes

https://github.com/Tijl/RSVP-repetitions-test

### Competing Interest Statement

The authors have declared no competing interest.

## References

Delorme, A., & Makeig, S. (2004). EEGLAB: An open source toolbox for analysis of single-trial EEG dynamics including independent component analysis. Journal of Neuroscience Methods, 134(1), 9–21. https://doi.org/10.1016/j.jneumeth.2003.10.009

Grootswagers, T., Robinson, A. K., & Carlson, T. A. (2019). The representational dynamics of visual objects in rapid serial visual processing streams. NeuroImage, 188, 668–679. https://doi.org/10.1016/j.neuroimage.2018.12.046

Grootswagers, T., Robinson, A. K., Shatek, S. M., & Carlson, T. A. (2021). The neural dynamics underlying prioritisation of task-relevant information. Neurons, Behavior, Data Analysis, and Theory, 5(1), 1–17. https://doi.org/10.51628/001c.21174

Grootswagers, T., Wardle, S. G., & Carlson, T. A. (2017). Decoding Dynamic Brain Patterns from Evoked Responses: A Tutorial on Multivariate Pattern Analysis Applied to Time Series Neuroimaging Data. Journal of Cognitive Neuroscience, 29(4), 677–697. https://doi.org/10.1162/jocn_a_01068

Grootswagers, T., Zhou, I., Robinson, A. K., Hebart, M. N., & Carlson, T. A. (2022). Human EEG recordings for 1,854 concepts presented in rapid serial visual presentation streams. Scientific Data, 9(1), 3. https://doi.org/10.1038/s41597-021-01102-7

King, J.-R., & Wyart, V. (2021). The Human Brain Encodes a Chronicle of Visual Events at Each Instant of Time Through the Multiplexing of Traveling Waves. Journal of Neuroscience, 41(34), 7224–7233. https://doi.org/10.1523/JNEUROSCI.2098-20.2021

Marti, S., & Dehaene, S. (2017). Discrete and continuous mechanisms of temporal selection in rapid visual streams. Nature Communications, 8(1), 1955. https://doi.org/10.1038/s41467-017-02079-x

Mohsenzadeh, Y., Qin, S., Cichy, R. M., & Pantazis, D. (2018). Ultra-Rapid serial visual presentation reveals dynamics of feedforward and feedback processes in the ventral visual pathway. ELife, 7, e36329. https://doi.org/10.7554/eLife.36329

Oosterhof, N. N., Connolly, A. C., & Haxby, J. V. (2016). CoSMoMVPA: Multi-Modal Multivariate Pattern Analysis of Neuroimaging Data in Matlab/GNU Octave. Frontiers in Neuroinformatics, 10. https://doi.org/10.3389/fninf.2016.00027

Robinson, A. K., Grootswagers, T., & Carlson, T. A. (2019). The influence of image masking on object representations during rapid serial visual presentation. NeuroImage, 197, 224–231. https://doi.org/10.1016/j.neuroimage.2019.04.050

Shatek, S. M., Robinson, A. K., Grootswagers, T., & Carlson, T. A. (2022). Capacity for movement is an organisational principle in object representations. NeuroImage, 261, 119517. https://doi.org/10.1016/j.neuroimage.2022.119517

Teichmann, L., Moerel, D., Baker, C., & Grootswagers, T. (2022). An empirically-driven guide on using Bayes Factors for M/EEG decoding. Aperture Neuro, 1(8), 1–10. https://www.doi.org/10.52294/82179f90-eeb9-4933-adbe-c2a454577289

